# Inhibition of eIF5A hypusination improves pancreatic islet transplantation outcomes by promoting ischemia tolerance

**DOI:** 10.1101/2025.11.24.689446

**Authors:** Hajar Ouahmi, Filippo Massa, Elisa Allart, Marc Cougnon, Matthieu Rouleau, Emmanuel Van Obberghen, Marina Shkreli, Antoine Sicard, Didier F. Pisani

**Affiliations:** Université Côte d’Azur, CNRS, LP2M, Nice, France; CHU Nice, Service de Néphrologie Dialyse et Transplantation, Hôpital Pasteur 2, Nice, France

## Abstract

Pancreatic islet transplantation provides substantial benefits for individuals with Type 1 Diabetes (T1D). However, the low yield of the procedure limits its therapeutic potential, as multiple injections from different donors are needed to provide sufficient islets for a single recipient. Many islets are lost during preparation and transplantation, primarily due to ischemia/reperfusion injuries. GC7 (N1-guanyl-1,7-diaminoheptane), an inhibitor of eIF5A hypusination, improves the resistance of various cells to ischemia/reperfusion. Our study therefore explored whether ex-situ conditioning of pancreatic islets with GC7 during the preparation process could serve as a strategy to enhance the outcomes of pancreatic islet transplantation.

Conditioning of mouse primary islets with GC7 at the digestion step of the pancreas significantly improved the procedure yield. Indeed, islets showed better survival at 24h and 72h after isolation and major protection against ischemia/reperfusion injury. Finally, the transplantation of a suboptimal number of islets under the renal capsule of hyperglycemic mice demonstrated superior functionality and survival of islets conditioned with GC7 compared to control islets.

Collectively, these findings demonstrate that conditioning pancreatic islets with GC7 enhances their resistance to ischemia-reperfusion injury and significantly improves the efficiency of pancreatic islet transplantation. Hence the use of GC7 appears as a promising strategy to win one of the major challenges currently faced in the field of Type 1 diabetes.

## 1. Introduction

Type 1 Diabetes (T1D) is an autoimmune disease characterized by pancreatic beta cells destruction resulting in insufficient insulin secretion. Islet allograft transplantation has progressively emerged as a promising alternative to insulin injection in patients with severe T1D or associated with advanced kidney disease [1–5]. In this approach, pancreatic islets isolated from brain-dead donors are percutaneously injected into the patient’s hepatic portal vein to restore the beta cell population [1]. This procedure has been shown in several clinical trials to improve glycemic control and quality of life [3, 4, 6–8], and is now used for patients displaying T1D with severe hypoglycemia or after kidney transplantation [9].

Notwithstanding this progress, the widespread use of islet transplantation is still hampered by a series of limitations. Because the metabolic benefits are dependent on functional islet potency and the number of transplanted islets that survive engraftment remains low, recipients need to receive two to three injections spaced a few months apart, isolated from 2 to 3 pancreas donors. This is a major hurdle both at the patient level (immunization) and from a public health perspective (shortage of donors) [10, 11]. The low yield of the procedure is due to significant islet loss resulting mainly from the oxidative stress occurring during the isolation process and after engraftment (ischemia/reperfusion injury and instant blood-mediated inflammatory reaction (IBMIR)). Oxygen is essential for beta cells due to their high mitochondrial oxidative metabolism [12], and ischemia therefore leads to their dysfunction and triggers inflammatory responses in vivo [6]. Insufficient oxygen supply to the islets results in the formation of a central hypoxic zone correlated to islet size and leading to necrosis. Similarly, hyperoxygenation can cause damage to the islets, although in this case, the damage is confined to their outer regions. [6]. In short, the oxygenation issue is a critical factor to address with high priority when developing strategies to enhance islet survival during all the transplantation phases [1, 6, 7].

Eukaryotic initiation factor 5A (eIF5A) is a protein translation modulator known to undergo hypusination which is generated from spermidine by the successive catalytic action of deoxyhypusine synthase (DHPS) and deoxyhypusine hydroxylase (DOHH) [13, 14]. Inhibition of eIF5A hypusination using a specific competitive inhibitor of DHPS (GC7: N1-guanyl-1,7-diaminoheptane) improves the resistance of mammalian tissues exposed to the noxious effects of ischemia/reperfusion [15–20]. This protection is secondary to a decrease in oxygen consumption due to a reversible shutdown of mitochondrial activity, with maintaining energy homeostasis through anaerobic glycolysis, as well as to a limited oxidative stress due to reactive oxygen species production (ROS) and endoplasmic reticulum (ER) stress [17, 19–21]. These findings have been extended to preclinical kidney transplantation models in pigs, where GC7 administration to the donor enhanced both early and long-term graft recovery [16, 17]. Recently, we demonstrated that treatment with GC7 protects INS-1 and MIN6 insulinoma cells against anoxia/reoxygenation by slowing down their metabolism (reduction in mitochondrial activity and oxidative stress), which significantly improved cell survival by more than 50% [22].

Given the essential role of oxygenation in the survival of pancreatic islets during isolation and transplantation, we sought to develop a procedure based on the use of GC7 to protect the islets during the isolation process and in the face of various stresses including ischemia/reperfusion. Our results demonstrated that the addition of GC7 specifically during the pancreas digestion step led to an improvement in the quantity and quality of the islets. It is important to note that these GC7-treated islets showed a significantly improved ability to control blood glucose levels in streptozotocin-induced diabetic mice compared to untreated islets.

## 2. Material and Methods

### 2.1. Reagents

When it is not specified, drugs, buffer solutions, fetal bovine serum (FBS) and culture reagents were from Sigma-Aldrich Chimie (*Saint-Quentin-Fallavier, France*). N-guanyl-1,7-diaminoheptane (GC7) was synthesized by AtlanChim Pharma (*Nantes, France*) according to the previously described method [23]. GC7 was used at the concentration of 30 µM which has been already validated to protect organs and cells from ischemia/reperfusion [15–21].

### 2.2. Islet Preparation

#### General procedure

The pancreas was harvested from freshly sacrificed (cervical dislocation) mice (BALB/c and C57Bl/6JRccHsd, Envigo*, Paris, France;* age and gender are indicated in figure legends), and then infused *ex vivo* with a solution of M199 (Sigma-Aldrich Chimie) supplemented by 5 mL of a mixture containing collagenase P (1000 u/mL, Roche Diagnostics, *Meylan, France*). Pancreas was digested for 15 min at 37°C, dispersed and passed through a tissue mesh (500 µm). Islets were isolated by elutriation protocol [24], handpicked and left O/N in retrieval medium (RPMI, 5.5 mM glucose, 10% v/v FBS).

#### Additional steps for GC7 supplementation

1/ Freshly harvested pancreas were preserved during 4h in University Wisconsin (UW) solution at 4°C with GC7 addition (30µM).

2/ Pancreas were infused with 5mL of a mixture of collagenase and GC7 (30 µM) and digested 15 min at 37°C.

3/ Pancreas were infused with 5mL of a mixture of collagenase and GC7 (30 µM), stored at 4°C for 45 min and then digested 15 min at 37°C.

4/ Islets were handpicked and left O/N in retrieval medium (RPMI, 5.5 mM glucose, 10% v/v FBS) containing or not GC7 (30µM).

For all the experiments, the same number of pancreases for each analyzed group was processed at the same time and in alternating order.

For anoxia, islets were maintained for 6h in an airtight chamber containing a 100% N_2_ atmosphere. Oxygen deprivation was controlled using an OXYBABY® apparatus (WITT, *Morsang sur Orge, France*). Reoxygenation was performed 2h under 5%/95% CO_2_/air atmosphere.

### 2.3. Transplantation Procedure in Diabetic Mice and Follow-Up

Mouse experiments have been performed in accordance with national animal welfare guidelines in the context of the authorized animal experimentation projects by the French Ministry (APAFIS#35008-2022011310408103). The number of animals was calculated to limit inter-individual variation and reach a statistical level. No mice were excluded after transplantation, and no blind analysis or randomization was performed. For the surgeries, one mouse from each group is transplanted at the same time and in alternating order, then housed together. Ten-week-old male BALB/c male mice were fasted for 4h and then treated with a single injection of streptozotocin (STZ, 200 mg/kg). Islet beta cell necrosis due to STZ toxicity was assessed by monitoring blood glucose levels the week after. Mice were transplanted when morning glycemia reached a level above 400 mg/dl.

For transplantation, a small incision was made on the left kidney, and a tunnel was created between the capsule and the cortex using a semi-rigid capillary (∼1 cm). Control or GC7 islets (∼150 IEQ = Islet EQuivalent to 150 islets with 150 µm of diameter), prepared following procedure described before using ten-week-old male BALB/c male mice, were then slowly delivered at the end of the tunnel using a tube mounted on a syringe (25 gauge-volume of 25 µL). The cut was then cauterized, and the kidney replaced in the cavity, the peritoneum and skin sutured (5.0 non-absorbable monofilament).

The capacity of the transplanted islets to reduce glycemia was assessed by following morning blood glucose for 4 weeks (10-11 a.m.) using glucometer (maximal value: 600 mg/dL) (Accu-chek, Roche Diagnostics, *Meylan, France*) and a drop of blood from the tail. An intraperitoneal glucose tolerance test (IP-GTT) was performed using O/N fasted mice at day 15 to evaluate the islet function. Mice were weighted and injected with glucose (2 mg/g body weight/500 µL NaCl 0.09%), and the glycemia monitored during 4h.

### 2.4. Glucose-stimulated insulin secretion (GSIS)

Islets were glucose-deprived for 1h in Krebs-Ringer Bicarbonate-Hepes buffer (KRBH) containing 1 mM glucose and 0.1% BSA (w/v). Then the cells were incubated for 1h30 in fresh KRBH containing 1- or 16-mM glucose. Centrifugated media and cell homogenates obtained by ethanol acid lysis were analyzed for insulin content using a mouse insulin ELISA kit (Mercodia AB, *Uppsala, Sweden*). Insulin secretion was expressed as the percentage of the cellular insulin content, which is referred to as fractional insulin release.

### 2.5. Immunofluorescence

Islets were stacked on Cell-Tak (Corning, Sigma Aldrich Chimie), fixed (paraformaldehyde 4%, 20 min, RT) and permeabilized (TritonX-100 0.3%/BSA 1%, 3h, RT) [25]. Islets were then incubated (O/N, 4°C) with CoraLite®488-conjugated anti-insulin (#CL488-66198, Proteintech, *Manchester, UK*) and CoraLite®594-conjugated anti-glucagon (#CL594-67286, Proteintech).

Islet-grafted kidneys and matched pancreas were sampled at the end of glycemia follow-up, fixed (paraformaldehyde 4%) and paraffin embedded. Sections (4 µm) were dewaxed and treated in boiling citrate buffer (10 mM, pH6.0, 15 min) for unmasking antigens. Cooled sections were rinsed and then permeabilized (0.2% TritonX-100, RT, 20 min), saturated with goat serum (3%, 30 min) and incubated (1h, RT) with CoraLite®488-conjugated anti-insulin, and/or CoraLite®594-conjugated anti-CD31 (#SAB5700639, Sigma-Aldrich and FlexAble 2.0 CoraLite® Plus 594 Antibody Labeling Kit KFA509, Proteintech). Slices were finally incubated with DAPI (#62248, Invitrogen) to visualize nuclei.

Finally, islets and slices were mounted with Histomount Solution (Thermo Fisher Scientific, *Illkirch, France*). Pictures were acquired with an Axiovert microscope (Carl Zeiss, *Le Pecq, France*) and processed with Zen software (Carl Zeiss).

### 2.6. Cell Survival Analysis

Viability of islets were evaluated by using Calcein-AM (#C3099, Thermo Fisher Scientific) and Ethidium homodimer (#E1903, Sigma-Aldrich Chimie). Fluorescent micrographs were analyzed with Fiji software using homemade macro [26]. Results are expressed as dead/live islet surface.

### 2.7. 3D islet culture

Matrix solution was prepared at 4°C in DMEM (5.5mM glucose, 1% FBS v/v) with PureCol® (1.7 mg/ml, Advanced BioMatrix, Thermo Fisher Scientific) and Matrigel Matrix® (25% vol/vol, Corning, Thermo Fisher Scientific) supplemented with HEPES 10 mM, glutamine 0.5 mM, 187.5 µL NaOH, 12 µL NaHCO3. A first matrix layer was placed in the well (400µL/well of 24-wells plate) and incubated at 37°C for two hours allowing matrix to gel. Ten pancreatic islets, isolated the day before, were mixed with 400 µL of the same cooled matrix solution and added to the top of the first matrix layer. Islets were also cultivated for 2 weeks in adequate medium (RPMI, 5.5 mM glucose, 10% v/v FBS) supplemented or not with 100 ng/mL of murine VEGF. Medium was changed every 72 – 96 hours.

After 5 days sprouting originated from the islet was visible. CD31 staining has confirmed endothelial phenotype of this sprouting which did not express insulin (Figure 1A). Neovascularization was evaluated using microscopic pictures of the pancreatic islets obtained by phase contrast at different times (from day 5 to day 14). Number of islets displaying sprouting and quantity of this vascularization were defined by measuring islet and sprouting areas as displayed in Figure 1B-C for CTRL islets isolated from BALB/c male mice (12-weeks old).

**Figure 1.**
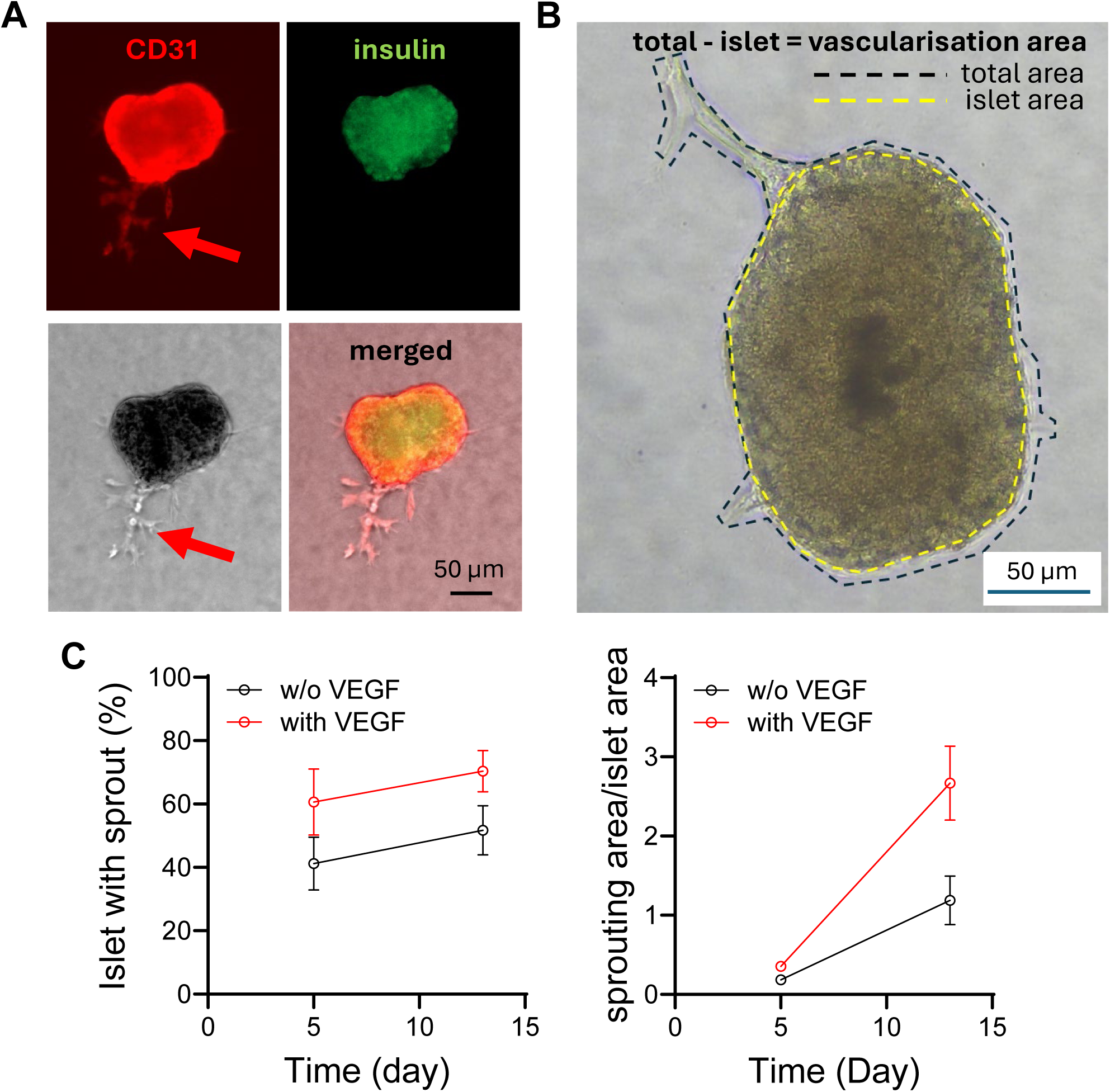
Islet culture to analyze neovascularization. slets isolated from the pancreas of BALB/c mice were cultivated for 13 days in matrix allowing vascular extension. (A) Microscopic picture of mouse pancreatic islet at day 9 in 3D culture, example of the insulin staining coupled with CD-31 staining to reveal vascular structure (including sprouting). (B) Microscopic picture of pancreatic islet presenting sprouting at day 9 of culture to measure the sprouting area according to the technic explained (subtraction of the core islet area of the total area to obtain the sprouting area). (C) Curves display the percentage of live islets presenting vascular extensions (n=4-12) and the mean area of these extensions normalized by islet area (n=30) after 5 or 13 days in the presence or not of exogenous VEGF.

### 2.8. Statistical Analysis

Results were analyzed by GraphPad Prism 7 software and using Mann–Whitney test to assess statistical differences between two experimental groups, or one or two-way ANOVA followed by Tukey’s post hoc test for multiple comparison. Differences were considered statistically significant with p<0.05. Number of replicates are indicated in the legend of figures.

## 3. Results

### 3.1. Development of a protocol for isolating pancreatic islets that enables the protective effects of GC7

To transpose GC7 protection to pancreatic islet preparation, we tested several protocols corresponding to procedure currently used in clinic, from donor pancreas harvesting to islet culture, using an adapted protocol with young male BALB/c mice. We first added GC7 (30 µM) to a UW solution (4°C, 4 h), a stage corresponding to the preservation of the pancreas after its removal from the donor. We then prepared the islets according to the standard protocol and, after a 24-hour recovery phase in a complete medium, we analyzed preparation with DTZ staining (Figure 2A), that allowed us to exclude residual exocrine tissue from the analysis, coupled to cell death analysis within the islets using a Live/Dead fluorescent test (Figure 2B). With this approach, GC7 added to the UW solution sensitized islet cells to the stress experienced during the procedure, as we found that GC7 strongly increased cell death (Figure 2C). The same type of results was observed for GC7 conditioning of already isolated islets. In this procedure, the pancreas was digested ex vivo immediately after being removed from the donor mouse, and the isolated islets were treated with 30 µM GC7 during the recovery phase (24 hours, 37°C, RPMI 10% FBS, 5.5 mM glucose). Analysis of pancreatic islet cell death at the end of the recovery phase or the following day, with or without GC7 remaining in the medium, showed an increase in mortality compared to untreated islets, which was further correlated with the duration of treatment (Figure 2C).

**Figure 2.**
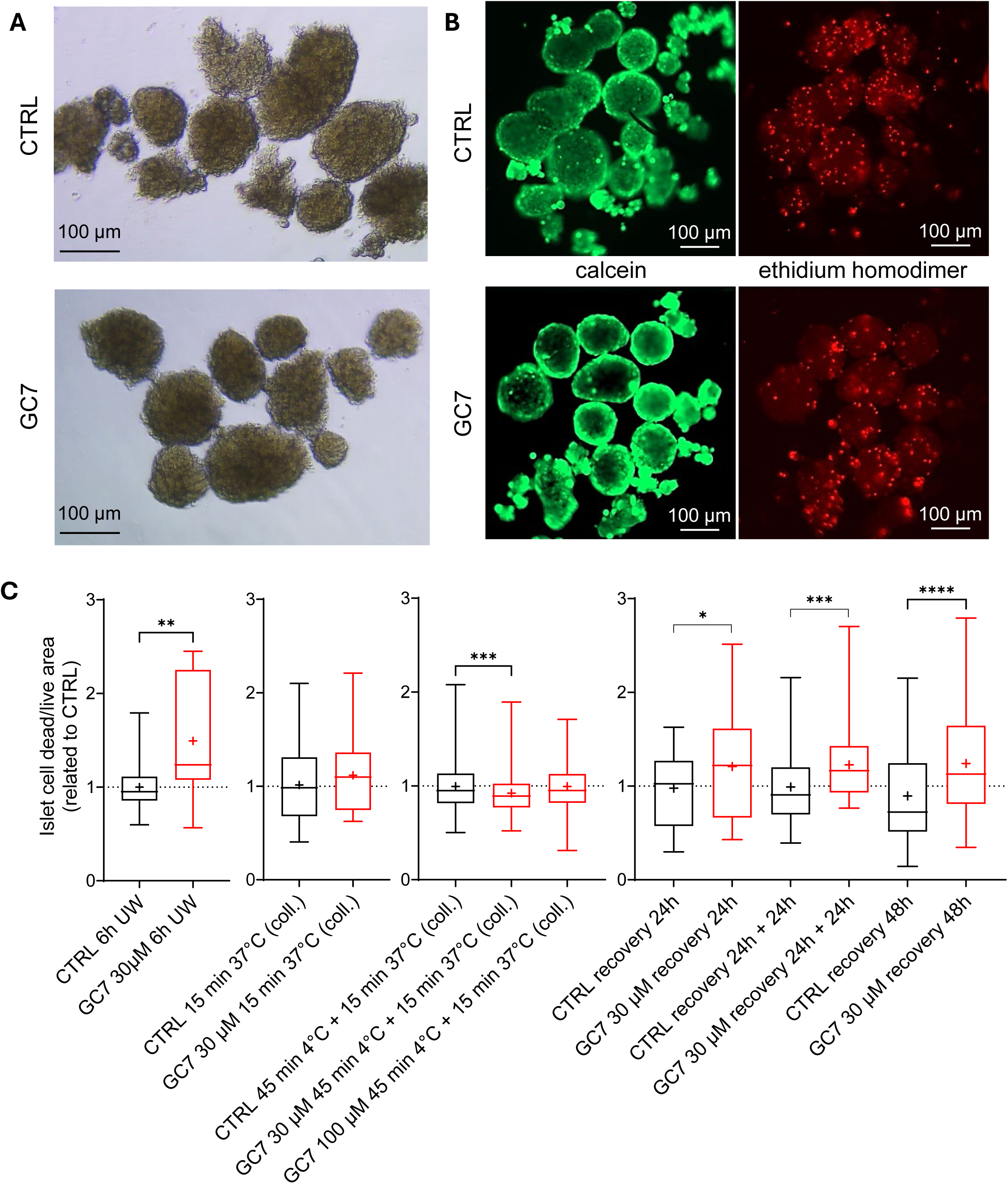
Effect of GC7 used at different step of pancreatic islet isolation procedure. BALB/c mouse (ten-week-old, male) pancreas was used to prepare islets following several procedures without or with GC7. (A) Example of islets at the end of recovery phase 24 h after isolation. (B) Islet cell survival was assessed by staining dead cells with Ethidium homodimer (red) and live cells with Calcein-AM (green). (C) Cell survival within islets according to various protocol of GC7 addition: addition to UW solution (4°C, 6h), during collagenase digestion with different time, temperature and concentration as indicated, and finally after isolation during the recovery phase in complete medium during 24 or 48h. Boxes show mean, value distributions and minimum and maximum values ((C) n=50-500, N=2-5). Statistical significance of mean differences was assessed by Mann-Whitney test (GC7 group vs. adequate CTRL group). * p<0.05, ** p<0.01, *** p<0.001, **** p<0.0001.

Finally, we defined a protocol consisting of a short treatment of the pancreas during the first stage of the *ex situ* digestion phase. In a first assay, we added 30 µM of GC7 to the collagenase mixture and directly digested the pancreas for 15 minutes at 37°C. Unlike previous procedures, under these conditions, treatment with GC7 did not alter islet cell mortality 24 hours after isolation (Figure 2C). Suspecting that the short treatment time was not sufficient to allow GC7 to take effect, we extended this time with a cold step. In this case, the procedure consisted of perfusing the ex vivo pancreas with a cold mixture of collagenase containing 30 µM of GC7 and leaving it in this medium for 45 minutes at 4°C, before activating digestion for 15 minutes at 37°C. Fortunately, this protocol improved the survival of pancreatic islet cells when prepared with GC7 compared to untreated cells (Figure 2C). This positive result could be due to sufficient but limited incubation time with GC7, as the drug added originally at 30 µM was only detected

in very low concentrations in the islet culture medium at the end of isolation (< 1 µM) and not at all at the end of the recovery phase. It should be noted that increasing the GC7 concentration to 100 µM using the same protocol negated its positive effect on cell survival (Figure 2C).

### 3.2. Pancreas per-conditioning with GC7 increases the yield and quality of pancreatic islets

To confirm the protective potential of GC7 when used during the prolonged cold digestion step, we thoroughly analyzed the quality of the islets 24 hours after their isolation. As shown in Figure 3A, dithizone staining, which is directly reflecting insulin content, was comparable in GC7-conditioned and control islets, and revealed islets with intact structures. This was confirmed by the immunofluorescence detection of insulin and glucagon in both groups (Figure 3A). Next, we evaluated the size and survival of a larger number of islets by analyzing the number of dead cells per islet. While the number of islets after the recovery phase was equivalent in both groups, we confirmed the improvement in cell survival per islet thanks to GC7 conditioning (Figure 3B). It should be noted that the treatment homogenized the size of the isolated islets (Figure 3C), demonstrating better structural preservation, and that the protective effect of GC7 was more pronounced in larger islets (Figure 3D).

**Figure 3.**
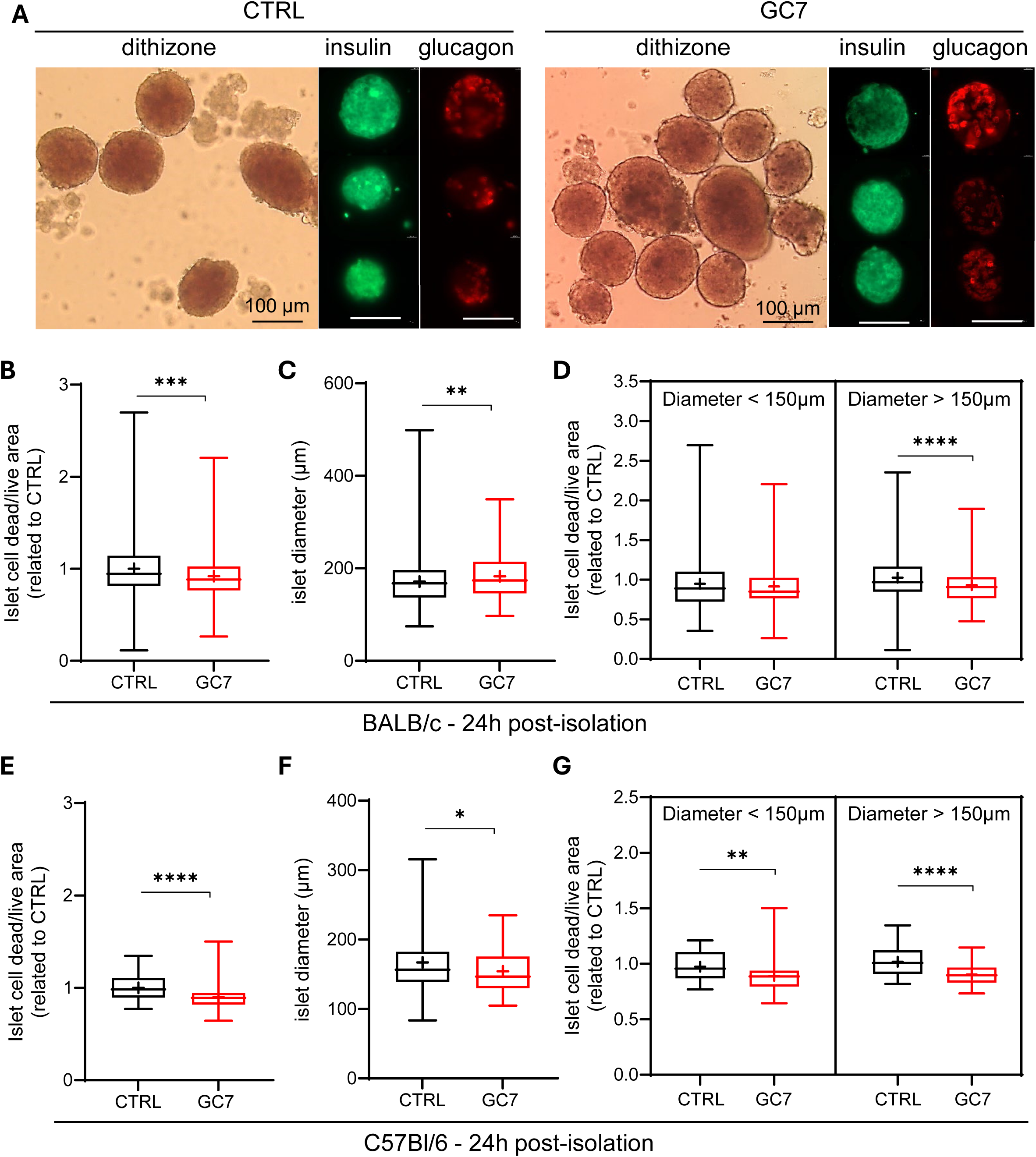
Effect of GC7 per-conditioning during pancreatic islet isolation. Mouse pancreas was used to prepare islets following the procedure without or with GC7 per-conditioning (30 µM). (B) Dithizone staining, insulin and glucagon immunofluorescence of islets 24h after isolation from BALB/c mouse pancreas. (B-G) The protective effect of GC7 per-conditioning was determined 24h after isolation from (B-D) BALB/c or (E-G) C57Bl/6 mouse (ten-week-old, male) pancreas by assessing (B,E) cell survival within islets, (C,F) islet diameter as well as (D,G) survival according to islet diameter. Boxes show mean, value distributions and minimum and maximum values ((B-D): n=70-120, N=3; (E-G): n=300, N=5). Statistical significance of mean differences was assessed by Mann-Whitney test. * p<0.05, ** p<0.01, *** p<0.001, **** p<0.0001.

We then evaluated the reproducibility of the procedure in another mouse strain using C57Bl/6 mice as pancreas donors (Figure 3E-G). Identically that for BALB/c mice, GC7 addition to the collagenase mixture and incubation at 4°C 45 min limited the mortality of pancreatic islet cells at the end of the recovery phase (Figure 3E), independently of their size (Figure 3G). Nevertheless, the size of the islets obtained with GC7 addition were again more homogeneous compared to control islets (Figure 3F).

### 3.3. GC7 enhances the survival of primary islets facing a stress

To mimic clinical conditions in which islets can be preserved for up to 3 days prior to transplantation, we cultured BALB/c pancreas-derived islets for 72 hours after the isolation procedure and evaluated their survival. First, evaluation at the end of the culture period of the number of islets with intact structure shows that while only ∼40% of the starting number of control islets were still intact and can be analyzed for cell survival, ∼60% of GC7-conditioned islets were preserved after 72 hours (Figure 4A). In line with the previous observation, the proportion of dead cells per viable islet was significantly lower in the GC7 group compared to the control one (Figure 4B). As shown in the right panels, the protection of large islets (i.e. above 150µm) was confirmed and is in fact more pronounced in prolonged cultivation conditions (Figure 4B).

**Figure 4.**
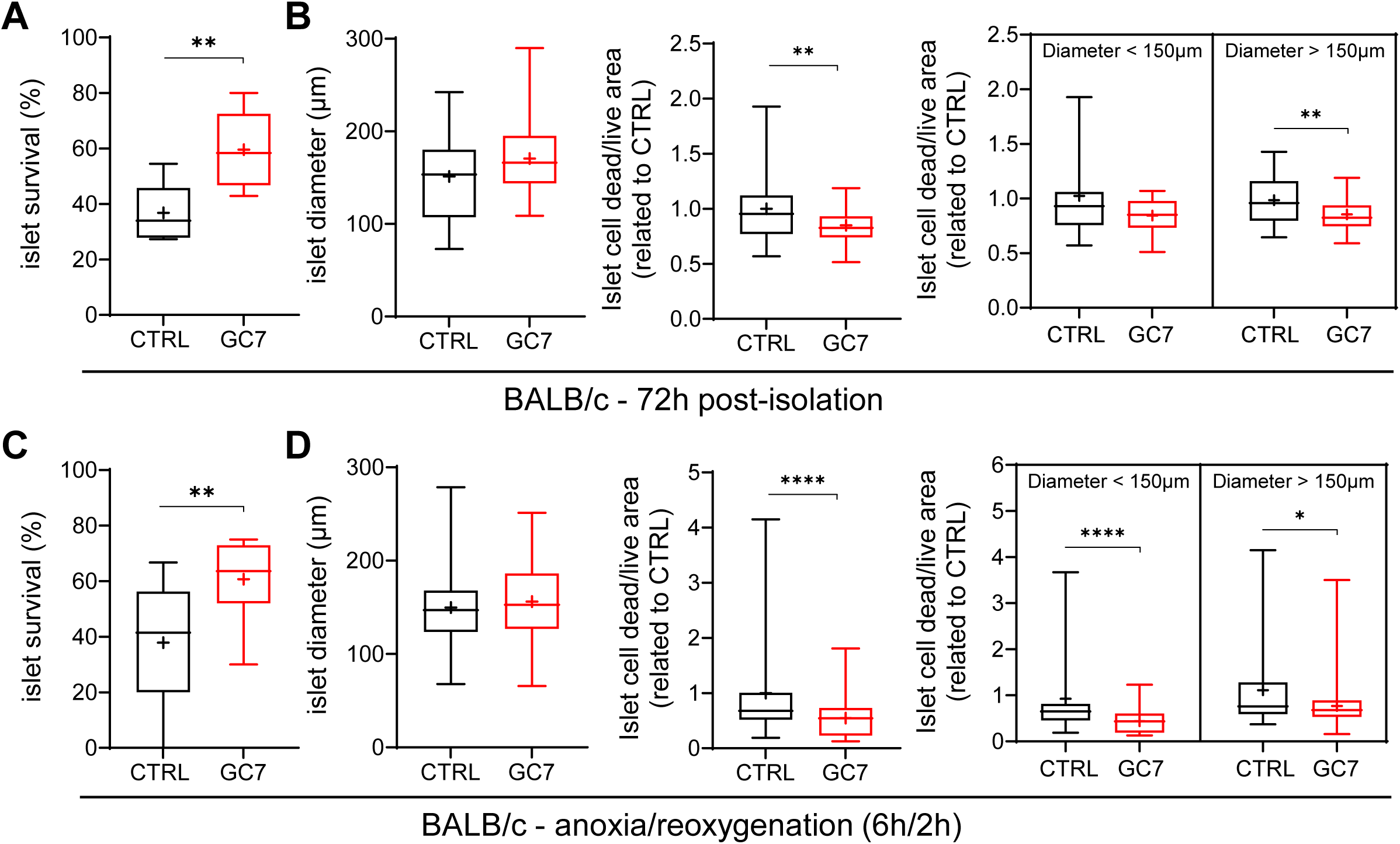
Effect of GC7 per-conditioning on islets submitted to stressed conditions. Islets were isolated from BALB/c mouse (ten-week-old, male) pancreas and, after 24h of recovery, (A-B) they were maintained for 48h more in culture or (C-D) they were submitted to anoxia/reoxygenation (6h/2h). (A, C) Islet survival, (B, D) diameter as well as cell survival within islets according or not to islet diameter were evaluated. Boxes show mean value distributions and minimum and maximum values ((A-B): n=80, N=3; (C-D): n=60, N=3). Statistical significance of mean differences was assessed by Mann-Whitney test. * p<0.05, ** p<0.01, **** p<0.0001.

Next, to evaluate resistance to the stressful conditions encountered during clinical islet transplantation, we exposed control and GC7-conditioned murine islets to anoxia/reoxygenation (6h/2h) after a 24-hour recovery period. Remarkably, while more than 60% of GC7-conditioned islets were still alive at the end of anoxia/reoxygenation, only about 40% of untreated islets were recovered (Figure 4C). The proportion of dead cells per viable islet was again significantly lower in the GC7 group independently of the islet diameter (Figure 4D). Unlike the results obtained at the end of the recovery phase, there is no homogenization of islet size obtained with the GC7 procedure after both stresses (Figure 4B, D). This is probably due to the loss of large islets in the CTRL group (values compared with Figure 3C), which are known to be more vulnerable, particularly to oxygen deprivation.

Islets derived from male and female C57Bl/6 mice displayed the same kind of resistance to anoxia/reoxygenation when they were prepared with GC7, with a decrease mortality of islet cells independently of islet sizes (Figure 5A,B).

**Figure 5.**
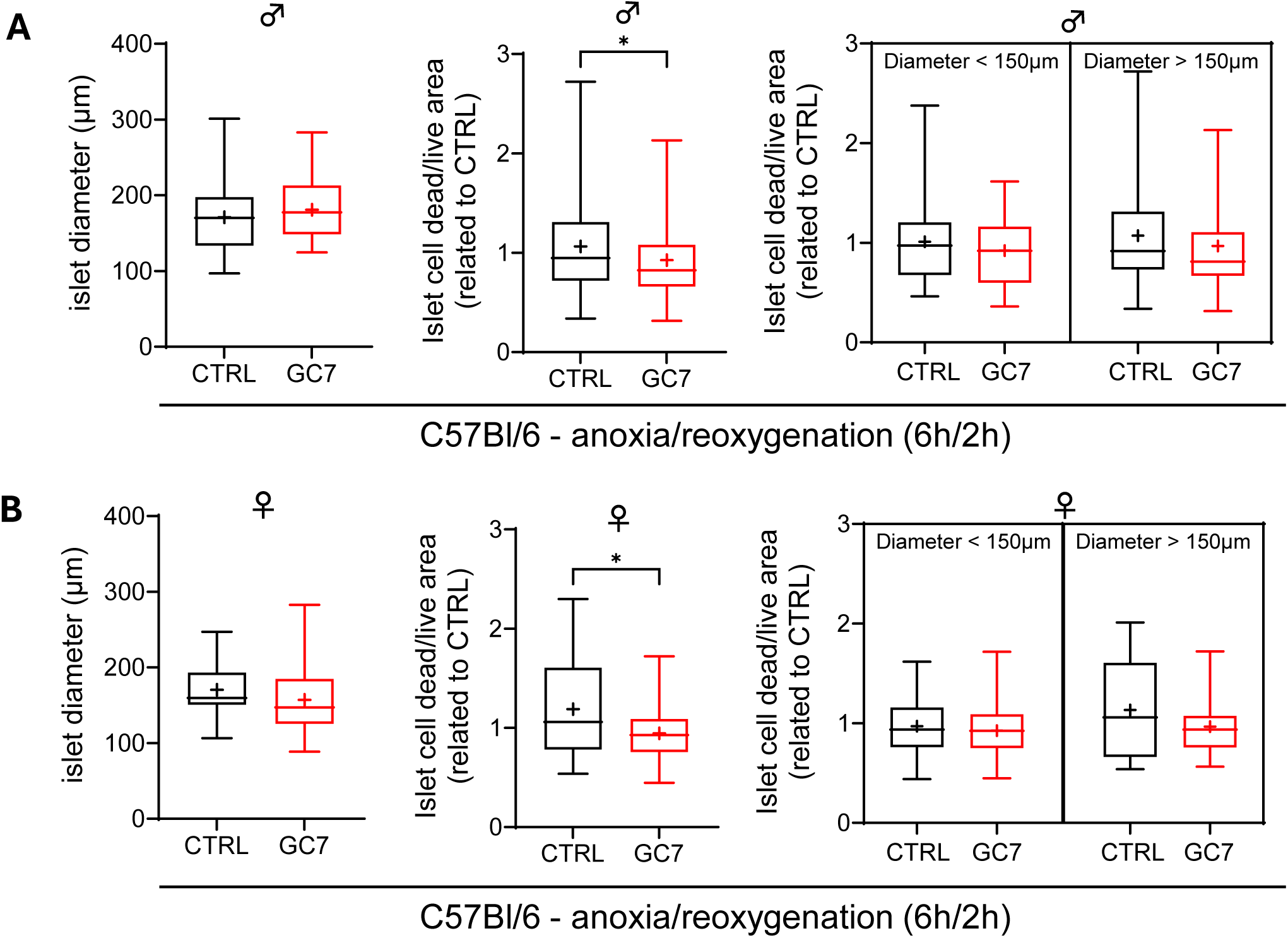
Effect of GC7 per-conditioning on islets submitted to anoxia/reoxygenation. Islets were isolated from male (A) or female (B) ten-week-old C57Bl/6 mouse pancreas and, after 24h of recovery, they were submitted to anoxia/reoxygenation (6h/2h). Islet survival, diameter as well as cell survival within islets according or not to islet diameter were evaluated. Boxes show mean value distributions and minimum and maximum values (n=15-90, N=3). Statistical significance of mean differences was assessed by Mann-Whitney test. * p<0.05.

### 3.4. GC7 protective effect was preserved with older donors

Human situation required the harvest of pancreas from donor with extended criteria including old donor. Taking account of this problematic, we prepared islets from 18-month-old female and male C57Bl/6 mice using our validated procedure. Analysis of survival after the recovery phase (Figure 6A-B) or in response to anoxia/reoxygenation stress (Figure 6C-D) showed that the protective effect of GC7 was maintained despite the age of the mice, but only for islets prepared from female mice (Figure 6A,C). Indeed, with islets prepared from aged male mice, preconditioning with GC7 resulted in a non-significant improvement in survival after 24 hours of recovery (Figure 6B), a trend that was confirmed after stress (Figure 6D) and independently of islet sizes. In old female mice GC7 preserved all islet independently of their size after the recovery phase, but it protected better bigger islets during anoxia/reoxygenation (Figure 6A,C). In old male mice, no significant protection was observed despite a downward trend in mortality among the largest islets (Figure 6B,D). Unlike young mice, we did not observe any homogenization of islet size in the GC7 groups compared to the CTRL groups (Figure 6A-D).

**Figure 6.**
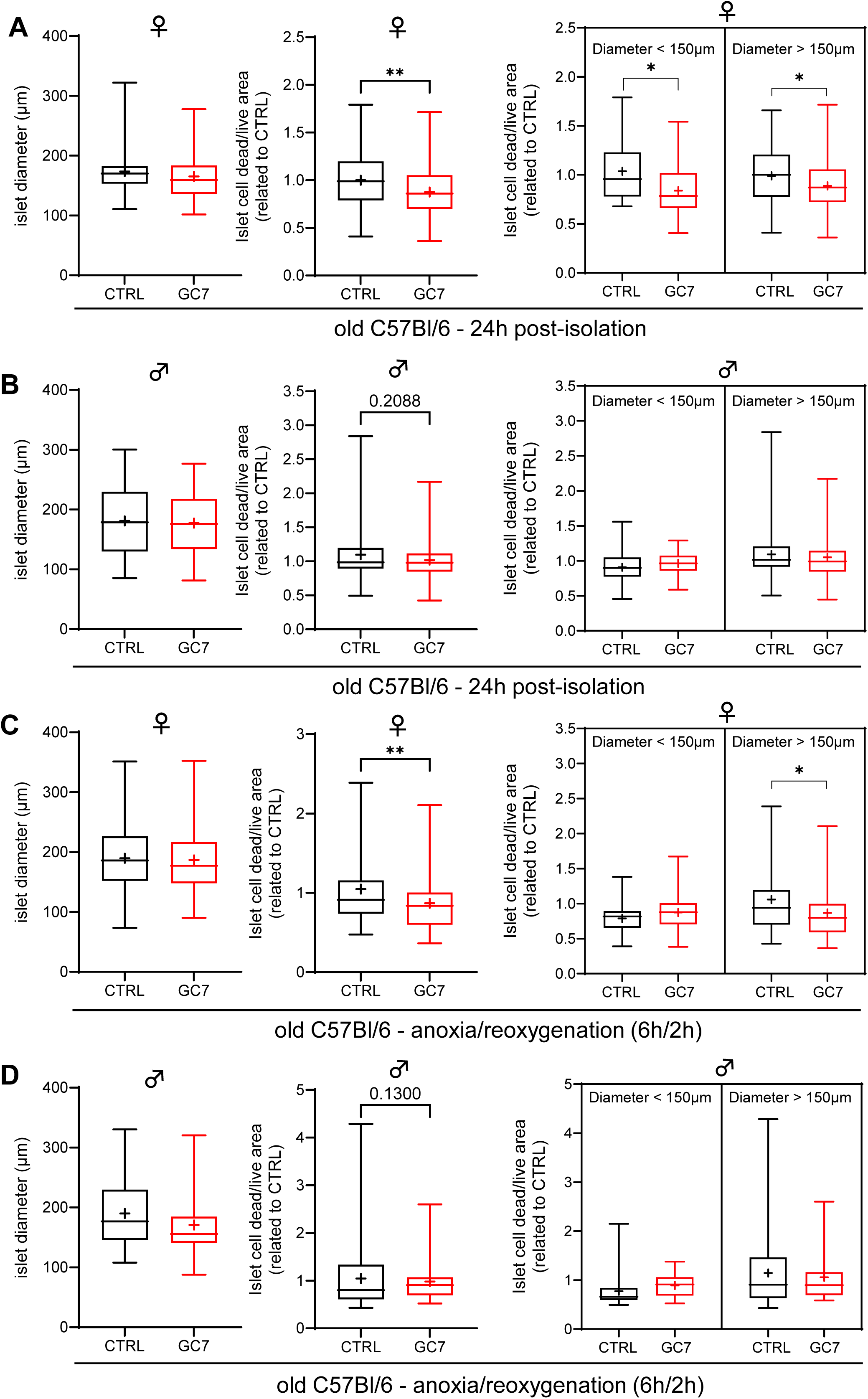
Impact of aging on GC7 protective effect. Islets were isolated from pancreas of 18-month old (A, C) female and (B, D) male C57Bl/6 mice after (A-B) 24h of recovery or (C-D) after additional anoxia/reoxygenation (6h/2h). (A-D) Islet diameter as well as cell survival within islets according or not to islet diameter were evaluated. Boxes show mean value distributions and minimum and maximum values (n=50-100, N=12). Statistical significance of mean differences was assessed by Mann-Whitney test. * p<0.05, ** p<0.01.

### 3.5. Islets neovascularization potential was affected by GC7 conditioning

Islet vascularization after transplantation is a crucial parameter for glucose sensing and systemic insulin action, and this can be achieved in part by the endothelial cells of the islet itself. Thus, to study the effect of GC7 on pancreatic islet neovascularization, we cultivated mouse islets prepared with or without GC7 in gel allowing sprouting of vascular tubes (Figure 7A). The vascular extension of pancreatic islets grown significantly across time in media with or without VEGF, and vascular area is higher in culture medium supplemented with VEGF compared to culture medium free of external VEGF (Figure 7B). Surprisingly, vascular area was always decreased when islets have been prepared with GC7 and cultivated without VEGF, a difference abolished in presence of exogenous VEGF (Figure 7B). Correlatively, the percentage of islets with neovascularization is significantly more important in control versus GC7 groups, respectively 35% to 1% in medium culture without VEGF (Figure 7C). This situation was normalized by the addition of VEGF which allowed around 50% of islets to vascularized in both groups (Figure 7C). Altogether, these results demonstrate that GC7 treatment blunts the capacity of the islets to promote its own vascularization by blocking VEGF production. A situation which could be counteract by recipient VEGF in vivo.

**Figure 7.**
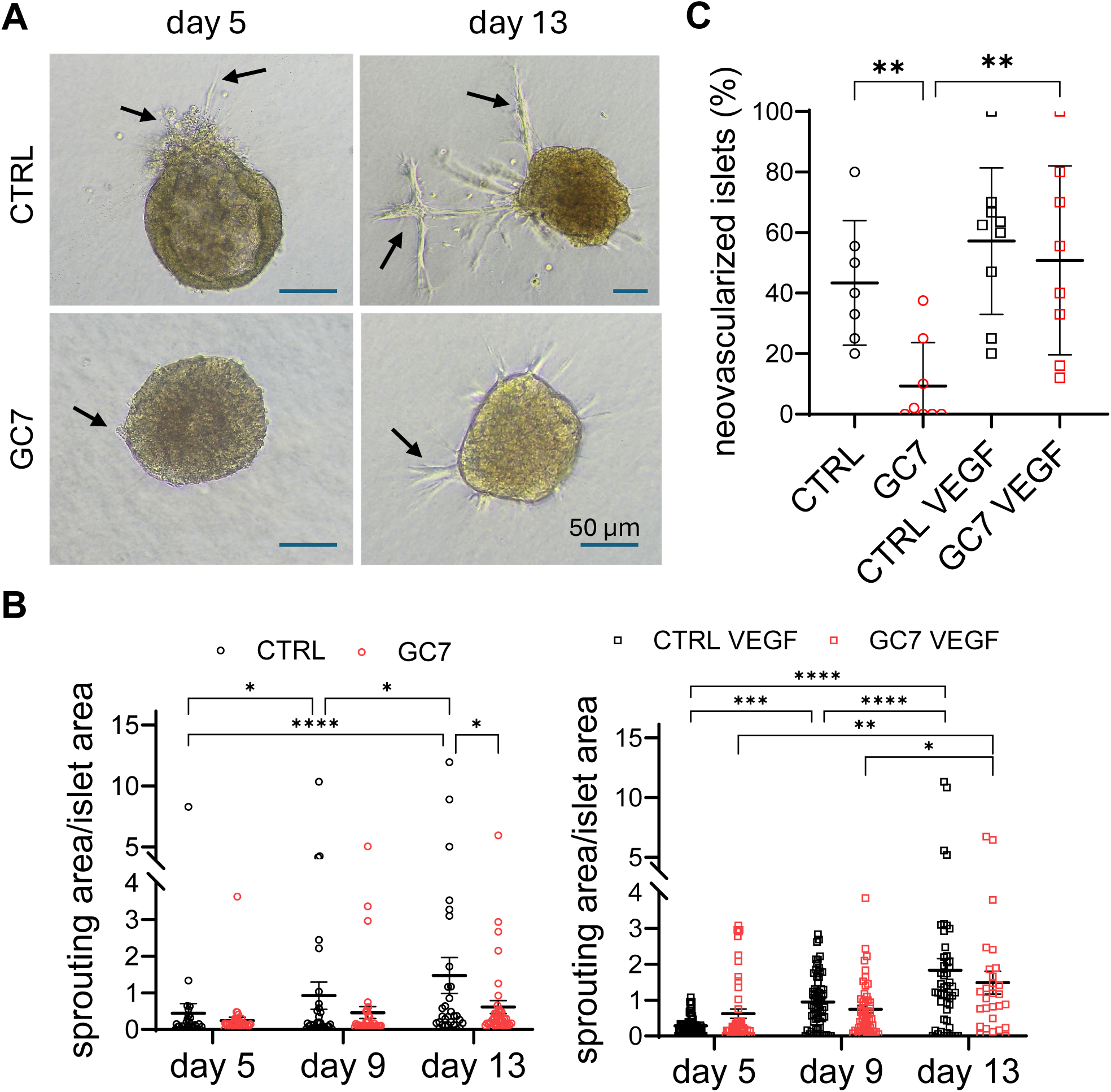
Effect of GC7 treatment at the time of pancreas processing on isolated islet neovascularization. Islets prepared with GC7 or not (CTRL) from the pancreas of BALB/c mice (ten-week-old, male) were cultivated for 13 days in matrix allowing vascular extension. (A) Example of neovascular network of CTRL and GC7 islets in 3D culture at day 5 and 13. (B) Vascular extension area normalized by islet area for CTRL or GC7 conditioned islets after 5, 9 or 13 days in the absence (left panel) or presence (right panel) of exogenous VEGF. (C) Percentage of live islets prepared with GC7 or not and presenting vascular extension after 13 days in the presence or not of exogenous VEGF (N=7-9). Dot plots display the individual values and mean +/- SEM (n=28-68, N=7-9). Statistical differences were assessed by one- or two-way Anova followed by Tukey post-hoc test. * p<0.05, ** p<0.01, **** p<0.0001.

### 3.6. Per-conditioned GC7 islets displayed higher therapeutic potential in diabetic mice

To reap the full benefits of the effects of GC7, we transplanted a sub-optimal quantity (∼150 IEQ) of untreated or GC7 conditioned syngeneic islets under the renal capsule of diabetic mice (mean glycaemia = 451 ± 26 mg/dl) (Figure 8A) and we assessed morning plasma glucose level for 3 weeks. Islet transplantation reduced glycemia in both groups of mice the day following surgery (Figure 8B). However, while mice transplanted with control islets failed to maintain their normoglycemia and quickly returned to the high blood glucose levels (> 400 mg/dL), the glycemia of mice grafted with GC7-conditioned islets continued to decline until day 5, stabilizing below 300 mg/dL until the end of the experiment at day 19. Measurement of overall glycemic variation showed that after 19 days, mice that received control islet transplants showed no improvement in glycemia compared to values observed before the transplant, while mice that received GC7-treated islet transplants showed a 20-50% improvement compared to their initial glycemic values (Figure 8C). Improved glycemic control was confirmed by IPGTT experiments demonstrating that mice transplanted with GC7-treated islets had a greater ability to respond to a glucose bolus by storing and utilizing it compared to the control group (Figure 8D).

**Figure 8.**
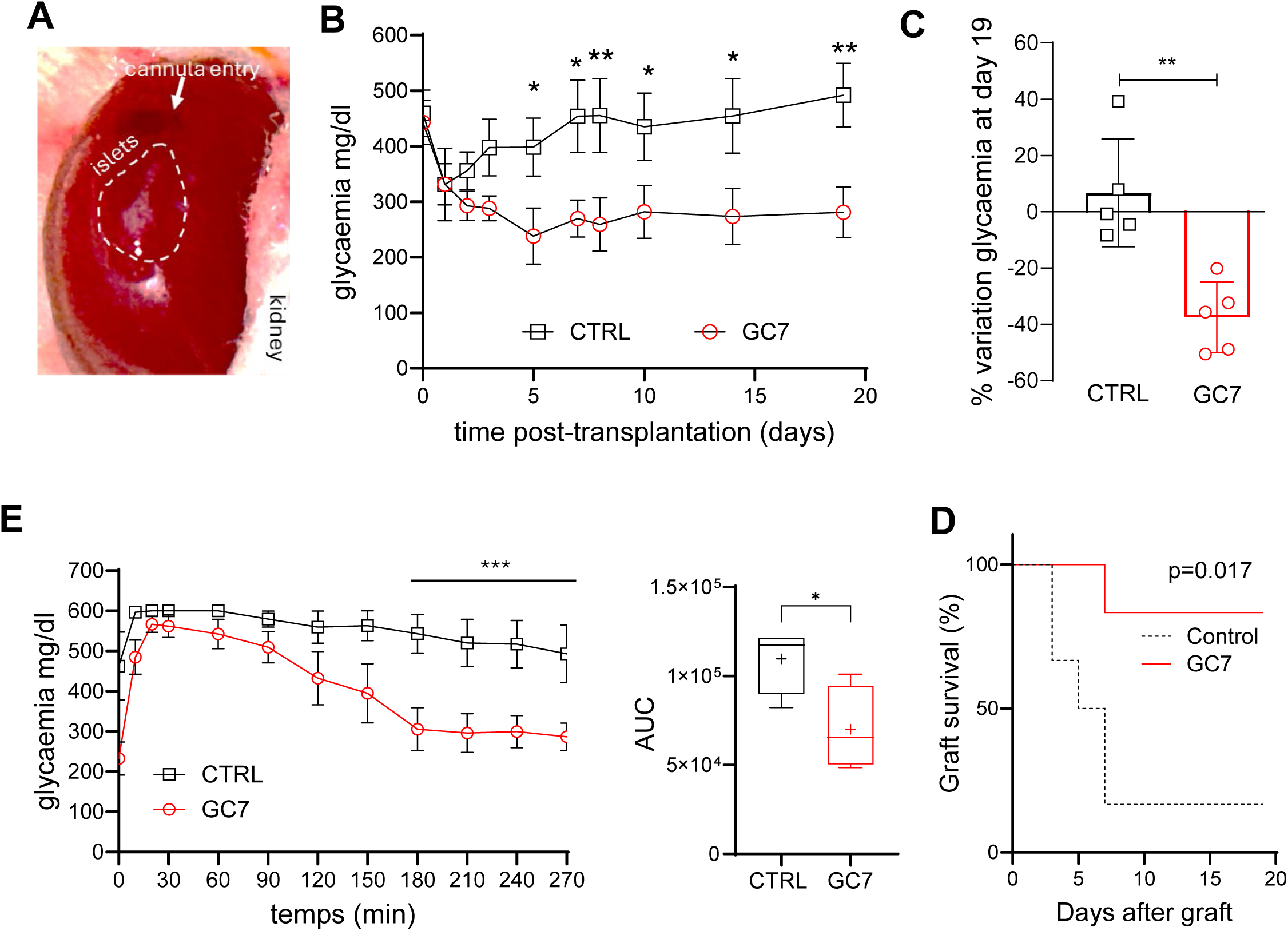
Therapeutic potential of GC7 per-conditioning islets for type 1 diabetes in mice. BALB/c male mice were injected with 200 mg/kg of streptozotocin. (A) 1-week later mice displaying a glycemia up to 450 mg/dL were transplanted with 150 IEQ untreated or per-conditioned with GC7 as shown in picture. (B) Glycemia was followed up to 19 days after transplantation (10:30 AM, fed mice). (C) % variation of glycemia at day 19 compared to pre-transplantation glycemia. (D) Curves displaying IP-GTT realized at day 15 post-graft (O/N fasted mice) and corresponding area under the curves (AUC). (E) Graft survival along follow-up. Mice with glycaemia higher than pre-transplant values are considered to have experienced “graft loss.” Curves and boxes show mean ± SEM, n=5. Statistical significance of mean differences was tested by two-way ANOVA followed by Tukey post-hoc test for curve and Mann-Whitney for comparison between two groups. * p<0.05, ** p<0.01, *** p<0.001.

Analysis of graft rejection/loss, characterized by post-transplant blood glucose levels higher than pre-transplant levels, showed that while 80% of mice that received a control pancreatic islet transplant lost their graft before day 10, 80% of mice transplanted with pancreatic islets prepared with GC7 retained their graft on day 19 (Figure 8E). This result was confirmed by immunohistological analysis of insulin expression in the transplanted kidneys and matched pancreas at the end of the follow-up period. While only few untreated islets were detected in the transplanted kidney area in control animals, islets prepared with GC7 were detected in quantity (Figure 9A, Figure 10A). To check if regeneration or preservation after STZ treatment of recipient pancreatic islets have participated to glycaemia normalization, we analyzed insulin expression in matched pancreas of transplanted mice. In order to verify whether regeneration or resistance to STZ treatment of the recipient’s pancreatic islets could have contributed to the normalization of blood glucose levels, we analyzed insulin expression in the matched pancreas of transplanted mice. As shown in Figure 9A, we found few or no islets in the pancreas of these mice, which rules out any involvement of this tissue in blood glucose control.

**Figure 9.**
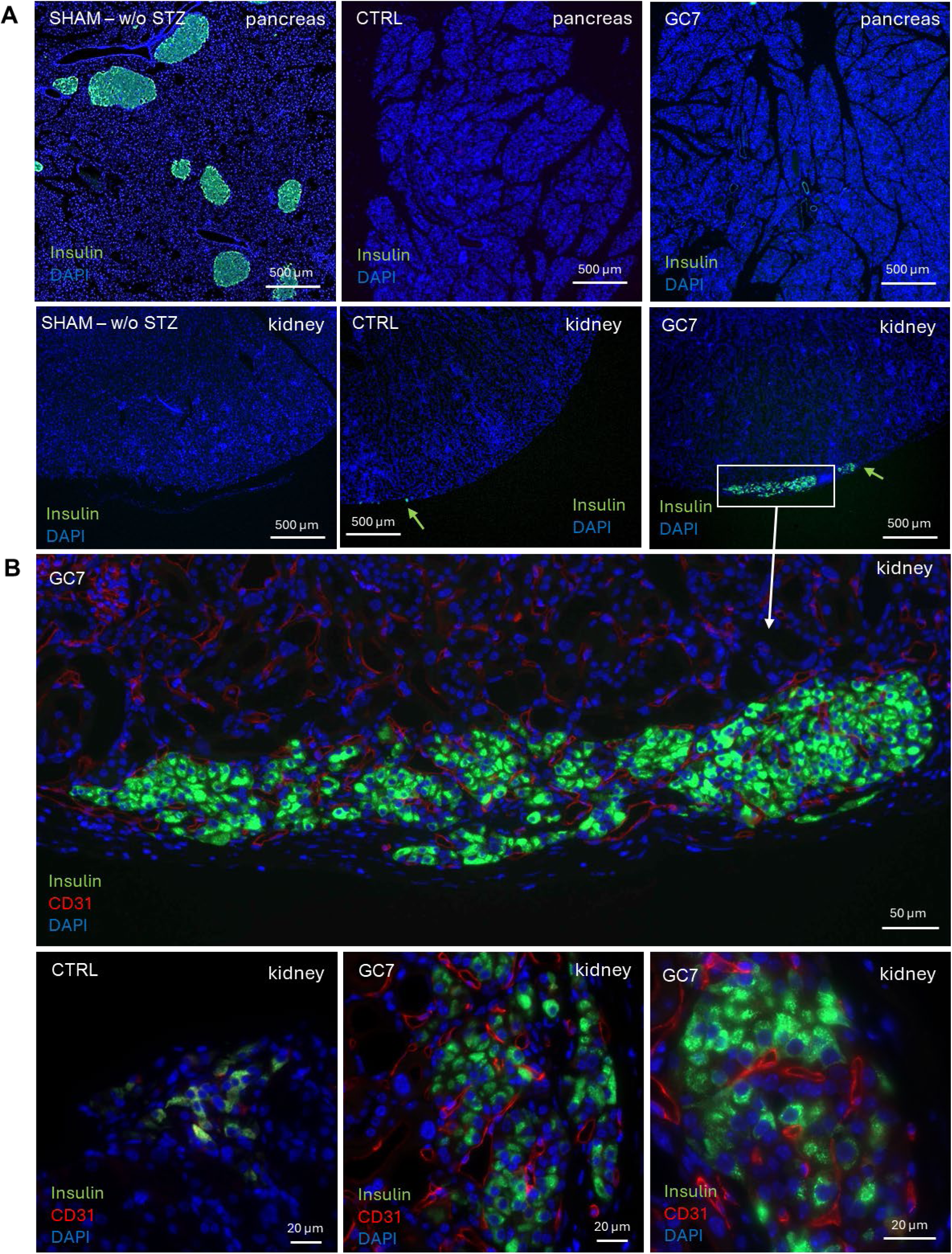
Characterization of GC7 per-conditioned islets after transplantation in DTZ-treated mice. BALB/c male mice were injected with 200 mg/kg of streptozotocin. 1-week later mice displaying a glycemia up to 450 mg/dL were transplanted with 150 IEQ untreated or per-conditioned with GC7. (A) Immunofluorescence analysis with anti-insulin antibody (in green) of renal subcapsular islet deposition sites and pancreas of the same mice at day 19 after transplantation. Pancreas and kidney of mouse untreated with STZ was displayed as control for staining and reference for normal islet content. (B) Magnification of islet deposition sites characterized by insulin positive cells (green) and vascularization by CD31 positive cells (red). (C) CD31 (red) and insulin (green) colocalization in the graft subcapsular kidney site 19-days after transplantation. DAPI was used to counterstained nuclei. Scale bars are indicated.

**Figure 10.**
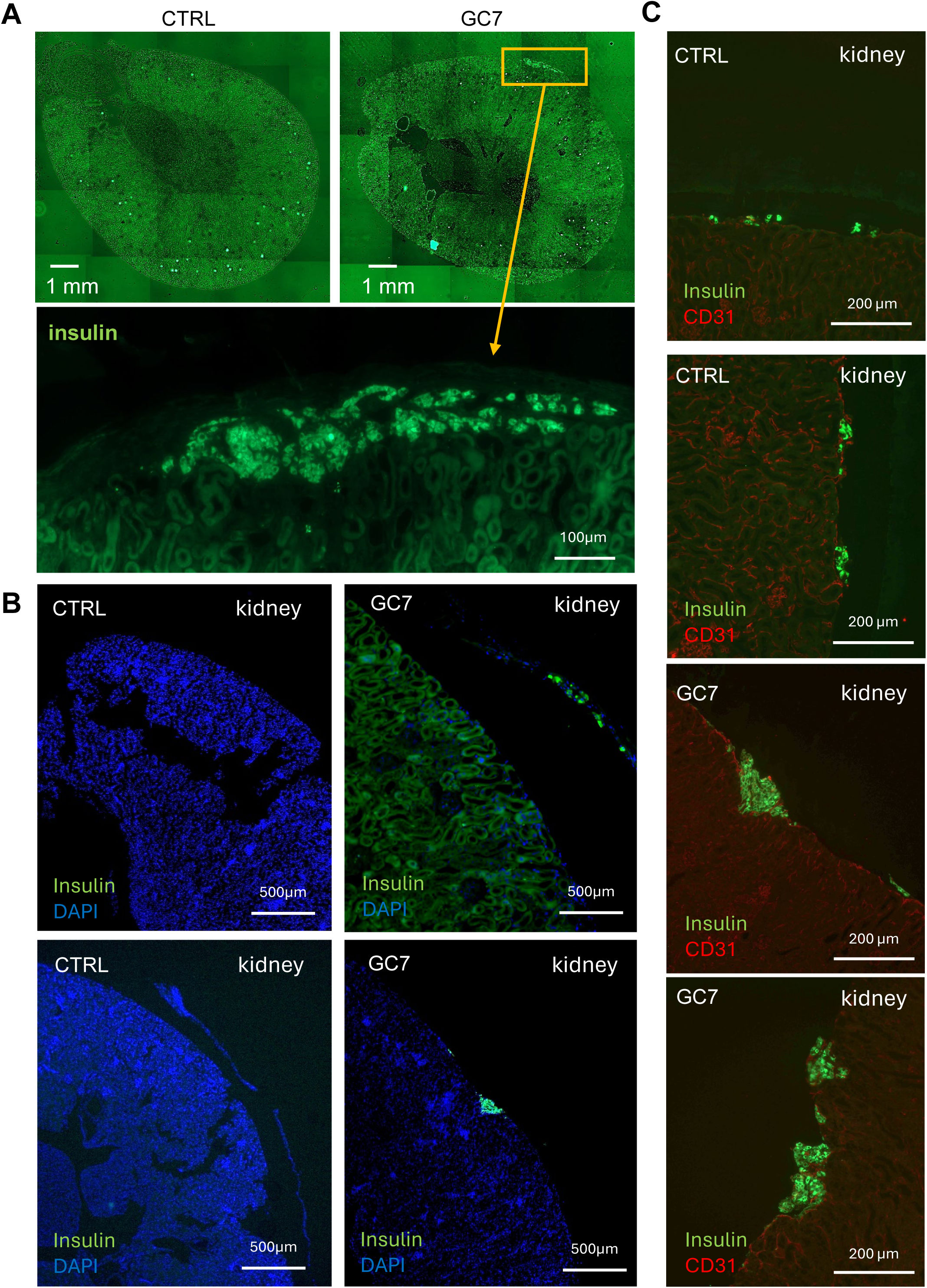
Characterization of GC7 per-conditioned islets after transplantation in DTZ-treated mice. BALB/c male mice were injected with 200 mg/kg of streptozotocin. 1-week later mice displaying a glycemia up to 450 mg/dL were transplanted with 150 IEQ untreated or per-conditioned with GC7. (A) Upper pictures display 5X mosaic of transplanted kidney with control or GC7-per-conditioned islets, and lower panel a magnification of the renal subcapsular zone containing islets. Only GC7 per-conditioned islets were retrieved on the graft site after 19 days. (C) Immunofluorescence detection of insulin (green) to detect islets in pancreas of DTZ-mice 19 days after subcapsular transplantation. (C) CD31 (red) and insulin (green) colocalization in kidney 19-days after transplantation. DAPI was used to counterstained nuclei. Scale bars are indicated.

Finally, the capacity to respond to blood glucose variation was permitted by a strong vascularization of GC7-treated transplanted islets under the renal capsule (Figure 9B, Figure 10B-C). This important result demonstrates that the inhibiting effect of GC7 on islet neovascularization found ex vivo unaffects the revascularization of grafted islets in the recipient.

Collectively, these data demonstrate that conditioning murine pancreatic islets with GC7 significantly improves their engraftment, maintenance and revascularization and thus the efficiency of islet transplantation.

## 4. Discussion

Type 1 diabetes (T1D) is due to selective destruction of beta cells, and its management is a most important public health challenge, due to the frequent morbid consequences of the disease [1, 27] with an increase incidence in young people last years [28–30]. The current challenges are to develop innovative strategies to support insulin secretion in situations where significant advances in insulin therapy have failed due to severe hypoglycemic episodes or long-term complications, as well as to improve the quality of life of patients with T1D by enabling them to no longer depend on exogenous insulin. Numerous studies and clinical trials demonstrate the efficacy and safety of pancreatic islet transplantation as a promising alternative for patients with severe T1D, improving patient outcomes and leading to insulin independence but limited in time. However, the beneficial effects of this approach are only achieved after two to three injections of beta cells spaced several months apart, requiring multiple donors for a single patient. This beta cell replacement therapy is therefore limited by the shortage of organ donors and the low yield of the pancreatic islet transplantation procedure. This is mainly due to graft rejection by the recipient, as well as insufficient vascularization and an unfavorable microenvironment in the final implantation site, contributing to ischemia/reperfusion-induced damage and thus beta cell suboptimal function and mortality [1, 7, 31]. Herein, we demonstrated that the use of GC7, an inhibitor of eIF5A hypusination known to be a highly effective anti-ischemic agent of great clinical interest in kidney transplantation and to protect rodent insulinoma cells from ischemia/reperfusion damage, can be of high interest to circumvent these problems.

Hypoxia during pancreatic islet transplantation is the primary non-immunological cause of cell death and graft failure, leading to persistent alterations in gene expression and impaired GSIS [32]. During the islet isolation process, oxidative stress resulting from oxygen deprivation during the isolation phase and subsequent reoxygenation during culture is a major contributor to islet injury. The native oxygen tension in pancreatic islets is approximately 40 mmHg, but in culture media conditions, oxygen levels are much higher (around 160 mmHg), which creates a significant discrepancy. Due to the loss of the islets’ native capillary network, oxygenation is largely dependent on diffusion, leading to insufficient oxygen delivery, especially to larger islets (>150 µm), which are prone to develop central necrosis [33–35]. In our study, GC7 preconditioning provided protection against ischemia/reperfusion injury during the entire isolation process, with a particularly beneficial effect on larger islets. This overall protection induced by GC7 improves the yield of transplantable islets, which is the key issue in islet transplantation [32]. After intraportal transplantation, islets lack a functional vascular system and have access only to the low oxygen level found in veins. Although islets acquire a functional capillary network within 2 weeks, oxygenation remains diffusion-dependent in the early post-transplant period [36, 37]. This oxygen problem could be more severe if the islets are transplanted in the subcutaneous space or intra-abdominally, two alternative grafting procedures suggested by recent papers [38, 39]. Various strategies, including islet encapsulation, have been developed to improve oxygenation [40–44], but these methods have not yet been successfully translated into clinical practice. Suszynski et al. identified four critical factors influencing islet oxygenation post-transplantation: (1) local oxygen partial pressure, (2) islet oxygen consumption, (3) islet size, and (4) thrombosis on the islet surface [32]. Interestingly, GC7 treatment reduced oxygen consumption in INS-1 cells, enhancing their resistance to hypoxemia by decreasing cellular oxygen demand. This relative tolerance to hypoxia decreased oxidative stress and improved cell survival. In vivo, GC7 preconditioning significantly improved glucose responsiveness in transplanted islets compared to untreated controls for up to 3 weeks. This period appears to be needed to reduce ROS and allow as such new capillary networks to form in the intraportal region [36, 37]. In our study histological analysis at the end of the follow-up revealed a loss of control islets, while GC7-preconditioned islets remained viable. These results concur with the loss of efficacy of the untreated islets to control glycemia on day 7 post-transplantation, which contrasts with a sustained reduction in glycemia of -35% on average up to day 19 post-transplantation of the GC7 treated islets. To sum up, these results support the notion that GC7 promotes islet survival by improving oxygen tolerance, particularly in larger islets, and by reducing oxygen consumption in beta cells.

The positive effect of GC7 on the resistance of islets and their effectiveness after transplantation could be offset by its inhibitory effect on the neovascularization of these same islets. Indeed, rapid revascularization is important for grafted pancreatic islets survival and function. It was demonstrated that transplanted islets are both revascularized by neovascularization from the islet himself and from the host vasculature [45]. Radial sprouting from the islet constitutes the angiogenic outgrow mediated by factors such as VEGF, it leads to the formation of vessel connections with nearby vessels and other islets. Vessels can sprout from the donor islet and interact with host cells involved in the vessel connection process [45]. Nevertheless, humoral immune responses significantly contribute to vascularized allograft failure due to donor specific antibodies primarily target the mismatched HLA molecules leading to antibody-mediated rejection that predominantly affects graft microvasculature [45]. However, in the context of pancreatic islet transplantation, as these are less vascularized, there is reduced possibility of this type of rejection, with revascularization originating mainly from the recipient [45]. Thus, inhibition of neovascularization from transplanted islets due to GC7 treatment could be considered as a positive effect in the context of pancreatic islet transplant rejection. This is particularly true given that neovascularization of the islets does not appear to be essential for the restoration of functionality in islets prepared with GC7, as these islets restore insulin secretion shortly after transplantation in diabetic mice.

The exact mechanisms enabling the GC7 to protect the islets must surely be very diverse. Polyamines, which are used as substrate for the hypusination of eIF5A, are essential for pancreas function especially for beta cells that displayed the higher content [46]. Consequently, polyamines depletion or chronic inhibition of hypusination impairs insulin secretion by beta cell function in response to glucose [47, 48] as well as beta cell mass [49]. Nevertheless, higher content of polyamines seems to be negative in pathologic conditions. Indeed, increase level of polyamines, and thus of hypusination, in beta cells correlated to T1D onset with an increase of endoplasmic reticulum (ER) stress and reactive oxygen species (ROS) production [50, 51]. It is interesting to note that another study has shown that polyamine depletion and inhibition of hypusination in mouse models of T1D may have a protective effect, preserving beta cell function and mass, and thus delaying the onset of the disease [52]. These findings are supported by previous results demonstrating that GC7 protects INS-1 and MIN6 insulinoma cells against ER stress [53, 54], and oxidative stress, the main mechanism of ischemia/reperfusion injury [22]. It is important to note that in the latter study, a correlation was established between the protective effect of GC7 and a slowing of mitochondrial activity and a decrease in oxidative phosphorylation, while simultaneously increasing anaerobic glycolysis in order to maintain adequate ATP synthesis for cell survival [22]. This divergence between physiological and pathological situations must also be considered in relation to the duration of inhibition. Indeed, even though the protective effect observed against ischemia/reperfusion is also associated with inhibition of cell function, the acute treatment used in these situations led to phenotypic consequences that were completely reversible after a short period of time [22] [21], thus limiting the adverse effects that prolonged inhibition of hypusination could have.

In conclusion, protecting islets from the detrimental effects of oxygen deprivation is essential for improving islet survival and function during both islet preparation and transplantation. Our findings presented here support the potential clinical application of GC7 as a conditioning agent to improve islet grafting outcomes. We believe that our approach could offer a unique opportunity for an improved diabetes life and health for the many young T1D patients waiting impatiently for disease modifying therapies.

## Acknowledgments

This work was supported by a grant from the Agence Nationale pour la Recherche (ANR-25-CE52-4257-02 MAXI-ISLETS) and Société Francophone du Diabète (2022). The authors greatly acknowledge Majlinda Topi and Maeva Bardin for technical assistance, and the IRCAN Animal Housing facilities and IRCAN Cytomed facilities that are supported by “le Cancéropôle PACA”, “la Région Provence Alpes-Côte d’Azur” and “le Conseil Départemental 06”. HO has a PhD fellowship “Espoir de la Recherche 2022“ from the Fondation pour la Recherche Médicale”.

## Data availability statement

All data used for analysis are available in the manuscript.

## Authors contribution

Antoine Sicard, Didier F. Pisani: Conceptualization, project administration, interpretation, planning administration, writing, funding acquisition, supervision, data acquisition and curation, methodology.

Hajar Ouahmi, Filippo Massa: Investigation, interpretation, data acquisition and curation, methodology, writing.

Elisa Allart, Marc Cougnon, Matthieu, Rouleau, Isabelle Rubera, Marina Shkreli: Investigation, data acquisition and curation, methodology.

Michel Tauc, Emmanuel Van Obberghen: Conceptualization, interpretation, writing.

